# Expansion of a single Helitron subfamily in *Hydractinia symbiolongicarpus* suggests a shared mechanism of cnidarian chromosomal extension

**DOI:** 10.1101/2024.11.21.624632

**Authors:** Tetsuo Kon, Koto Kon-Nanjo, Oleg Simakov

## Abstract

Helitrons are rolling-circle transposons that amplify through rolling-circle replication mechanism. Since Helitrons were relatively recently identified, their impact on genome evolution is still not fully understood. Here, we describe that a single Helitron subfamily specifically accumulates in the subtelomeric regions of *Hydractinia symbiolongicarpus*, a colonial hydrozoan cnidarian. Based on the sequence divergence, it is suggested that the Helitron subfamily underwent a burst of activity in the species’ recent history. Additionally, there is a IS3EU DNA element accumulation at the putative centromeric regions, as well as minisatellite sequences of approximately 200 bp in length extending from the telomere-side end of the Helitron towards the telomere. Phylogenetic analysis of Helitrons in the *H. symbiolongicarpus* genome suggests that the Helitrons underwent local propagation at the subtelomeric regions. The single Helitron subfamily, along with the consecutive minisatellite, accounts for 26.1% of the genome coverage (126 Mb of the 483 Mb genome), which collectively contribute to the genome size increase observed in *H. symbiolongicarpus* compared with other cnidarians. Homologous sequences of Helitron in *H. symbiolongicarpus* were identified in the genomes of other cnidarians, suggesting that Helitrons in hydractinia were present in at least the common ancestor of Cnidaria. Furthermore, in *Nematostella vectensis*, an anthozoan cnidarian, Helitrons were also accumulated at the subtelomeric regions. All these findings suggest that Helitrons constitute a common cnidarian mechanism of chromosomal extension through local amplification in subtelomeric regions, driving diverse genome expansions within the clade.

## Introduction

The genomes of animals, even after hundreds of millions of years of divergence, retain conserved sets of orthologs between species, which form ancestral linkage groups and maintain chromosomal-level homology [1,2]. In addition to protein-coding genes, transposable elements (TEs) are another major genomic elements that constitute the majority of noncoding regions in many animal genomes [3]. TEs are broadly classified into two major categories: RNA transposons (class I TEs, also known as retrotransposons), which transpose via an RNA intermediate through a copy-and-paste mechanism, and DNA transposons, which move through a DNA-based cut-and-paste mechanism [4]. As mobile elements, TEs can “jump” throughout the genome, exerting wide-ranging effects [4–6]. For example, TEs can insert into gene regulatory elements, altering gene expression, and thereby potentially contributing to the evolution of genome function in various ways [5]. Additionally, changes in the copy number of TEs can lead to changes in genome size, an important aspect of animal genome evolution. Dozens of TE families are constitutively found in the genomes of a wide range of animals (Active TEs, A-TEs) and are involved in genome size expansion [3]. Moreover, alongside these A-TEs conserved across animals, certain TE families exhibit particularly high activity in specific animal clades and may play a significant role in shaping the unique genome evolution of those species or lineages [7,8].

Helitrons are rolling-circle (RC) transposons in the major TE groups that were identified relatively recently, with their discovery in 2001 [9], over 50 years after McClintock’s groundbreaking identification of the first transposable elements, the Ac/Ds elements, in maize [10]. Among the various TEs, Helitrons are particularly notable for their unique mechanism of replication, which involves the rolling-circle replication process. In this process, a nick is formed in the genomic DNA, serving as the starting point for the generation of circular DNA [11–13]. This circular DNA is then inserted into regions of the genome [11–13]. Helitrons are classified into two functional types: autonomous and non-autonomous. Autonomous Helitrons encode the essential proteins necessary for their replication and transposition [12]. These proteins facilitate the single-strand DNA nicking and rolling-circle replication necessary for Helitron activity. In contrast, non-autonomous Helitrons lack these protein-coding regions, making them incapable of independent transposition. Instead, they depend on the enzymatic machinery provided by autonomous Helitrons for their amplification and mobility. Most of the Helitrons identified to date are nonautonomous [12]. Helitrons have been described in a wide range of animals and plants [14–19], and their deep evolutionary association with telomeres has been proposed [20]. However, the details of their replication mechanisms, diversity, and interactions with host genomes remain to be fully elucidated.

*Hydractinia symbiolongicarpus* is a colonial hydrozoan cnidarian species found along the east coast of North America, forming colonies on hermit crab shells and can be cultured in laboratories [21]. This species has specialized zooids (polyps) for feeding (gastrozooid), reproduction (gonozooid), and defense (dactylozooid), and its daily spawning of gametes facilitates genetic manipulation techniques like RNAi, CRISPR/Cas9, and transgenesis [22–28]. Notably, *H. symbiolongicarpus* was the first animal in which stem and germ cells were described, possessing unique pluripotent interstitial stem cells (i-cells) that can differentiate into all somatic and germ cell lineages [29]. The ability of *H. symbiolongicarpus i-cells* to differentiate into all cell types differs from the more limited potential of *Hydra vulgaris* i-cells, which can generate neuroglandular cells and germ cells but are unable to produce epidermal and gastrodermal epithelial cells [30–33]. Therefore, *H. symbiolongicarpus* is an attractive model for studying stem cell function and evolution, and recent studies using this species as a model have led to various significant biological discoveries [23,29,34–36].

In the previous study, we assembled the all 15 chromosome sequences of *H. symbiolongicarpus* by combining long-read sequencing with DNase Hi-C [37]. The genome of *H. symbiolongicarpus* retained chromosomal scale synteny with other Cnidaria, although colinear synteny was almost lost compared to *H. vulgaris*, which belongs to the same Hydrozoa class because of intrachromosomal rearrangements [3,37]. The genome assembly size was 483 Mb, more than 100 Mb larger than the genome sizes of many other Cnidaria, which typically range from 200-300 Mb [38–40]. Although 61% of the entire genome consisted of repetitive elements, most of them could not be attributed to specific TE families [37]. However, the distribution of sequence divergence in the repetitive elements suggested that there was a past burst of TE activity during a narrow time window [37].

The nucleotide sequences of TEs generally undergo rapid substitutions, and inactive TEs progressively becomes fragmented, making it often challenging to accurately annotate them to known TE families [41,42]. This tendency is particularly evident in animals other than well-studied model organisms such as mice and zebrafish, which have well-characterized genomes [3]. In this study, we used a highly sensitive TE analyses, based on the hidden Markov model (HMM) that we previously developed. Our findings suggest that the past TE burst in *H. symbiolongicarpus* was primarily driven by Helitrons. Furthermore, we also report that by utilizing genome coordinate information obtained from our previous chromosomal-scale genome assembly [37], we discovered that a single subfamily of Helitrons preferentially is accumulated in subtelomeric regions.

## Results

### TE coverage analysis in the *H. symbiolongicarpus* genome

In the genome of *H. symbiolongicarpus*, it was considered difficult to attribute the past TE bursts to appropriate TE families due to nucleotide substitutions and fragmentation of the TE remnants in the previous study [37]. Therefore, in this study, we created a custom repetitive element library using RepeatModeler [43] with modified method [3]. This modified method utilized an HMM-based sequence search [44], instead of using rmblast, which is employed by RepeatClassifier, a subprogram of RepeatModeler, to compare the sequences of the custom repetitive element library generated by RepeatModeler against the Dfam database [3,43]. In the original default run of RepeatModeler, 2,300 sequences were generated as a custom repetitive element library, of which 24.4% (562 sequences) were attributed to one of the TE families. On the other hand, the modified method provided improved annotation; we were eventually able to annotate 63.5% (1,461 sequences) of the total sequences. Using this refined custom repetitive element library as an input for RepeatMasker, we examined the TE landscape in the chromosome-level genome assembly of *H. symbiolongicarpus* (total size 483 Mb) [37]. As a result, we found that 38.0% (184 Mb) of the genome is occupied by SINEs, LINEs, LTRs, DNA, and RC elements (Table S1). When unclassified repetitive elements were also included, the genome coverage expanded to 60.4% (292 Mb). Specifically, SINEs accounted for 1.43% (6.93 Mb, compared to 0.350% [1.69 Mb] in the default RepeatModeler/RepeatMasker run), LINEs for 4.73% (22.8 Mb, compared to 2.93% [14.2 Mb] in the default RepeatModeler/RepeatMasker run), LTRs for 2.98% (14.4 Mb, compared to 1.77% [8.53 Mb] in the default RepeatModeler/RepeatMasker run), DNA elements for 17.1% (82.4 Mb, compared to 4.78% [23.1 Mb] in the default RepeatModeler/RepeatMasker run), and RC elements for 11.8% (57.1 Mb, compared to 0.561% [2.71 Mb] in the default RepeatModeler/RepeatMasker run) (Table S1). Furthermore, analysis of the nucleotide substitution distribution of TEs (Kimura divergence scores) revealed a sharp peak, as we previously reported [37], suggesting a past burst of TE activities (Fig. 1a,b). Although the main components of the sharp peak were unclear when using the default RepeatModeler custom repetitive element library (Fig. 1a), the analysis with the refined custom repetitive element library showed that Helitrons was main component in the sharp peak (Fig. 1b). This suggests that the *H. symbiolongicarpus* genome experienced a Helitron burst in the past.

**Fig. 1.**
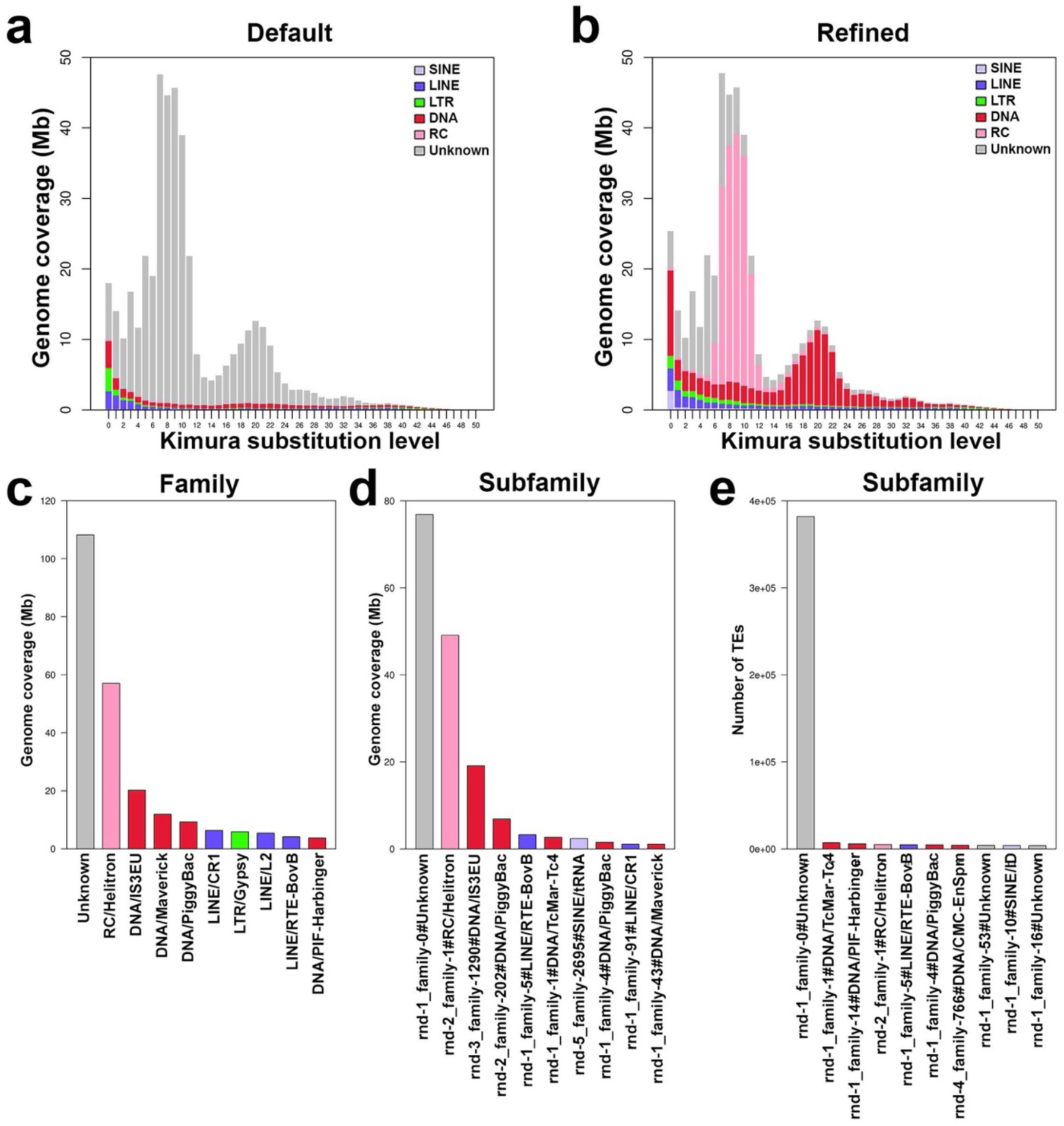
TE genome coverage in *H. symbiolongicarpus* genome. **(a)** Repeat landscape illustrating TE accumulation history for the *H. symbiolongicarpus* genome with the custom repetitive element library generated by RepeatModeler run with the default settings. The horizontal axis illustrates the Kimura substitution levels of repeat elements relative to their consensus sequences, indicating their ages. On the vertical axis, the graph depicts the genome coverage of each repeat family within the genome in Mb. Consequently, older repetitive elements appear toward the right side of the graph, while more recently active repetitive elements are positioned on the left. (**b**) Repeat landscape with the refined custom repetitive element library. (**c**) Genome coverage of TE families. (**d**) Genome coverage of TE subfamilies. (**e**) Copy number of TE subfamilies.

We examined the genome coverage of TEs at both the family and subfamily levels. At the family level, the most abundant elements were unclassified elements (22.4%, 108 Mb), Helitrons (11.8%, 57.1 Mb), and DNA element of IS3EU, (4.19%, 20.2 Mb) (Fig. 1c, Table S2). At the subfamily level, single subfamily of unclassified element (ID: rnd-1_family-0) occupied 15.9% (76.9 Mb) of the genome (Fig. 1d, Table S3), with an overwhelming copy number of 382,119 copies, far surpassing all other TE subfamilies (Fig. 1e, Table S3). This unclassified element (ID: rnd-1_family-0) represented 71.0% of all unclassified elements (Table S2,3). A single Helitron subfamily (ID: rnd-2_family-1) accounted for 10.2% (49.1 Mb) of the genome (Fig. 1d, S1a), representing 86.1% of all Helitron elements (Table S2,3). The copy number of the Helitron subfamily (ID: rnd-2_family-1) was 4,938 (Table S3). A single IS3EU subfamily (rnd-3_family-1290) accounted for 3.96% (19.1 Mb) of the genome (Fig. 1d, S1b), representing 94.4% of all IS3EU elements (Table S2,3). The copy number of the IS3EU subfamily (rnd-3_family-1290) was 1,384 (Table S3). The Kimura substitution level of this Helitron subfamily was 8.86%, which is the similar value of the RC peak (Fig. 1b, Table S3). Moreover, the Kimura substitution level for the subfamily of unclassified element (rnd-1_family-0) was 5.85% (Fig. 1b, Table S3). The single subfamily of IS3EU showed the Kimura substitution level of 20.6%, which corresponds to the second wide peak (Fig. 1b, Table S3). In summary, we identified an Helitron subfamily (ID: rnd-2_family-1, 10.2% of genome coverage, 49.1 Mb), an IS3EU subfamily (rnd-3_family-1290, 3.96% of genome coverage, 19.1 Mb), and an unclassified element (rnd-1_family-0, 15.9% of genome coverage, 76.9 Mb) that together account for 30.0% (145 Mb) of the genome. These elements are suggested to be associated with the genome size increase in the *H. symbiolongicarpus* genome compared with other cnidarians [38–40].

## Distribution of Helitron copies on chromosomes

Next, we examined how the subfamily of Helitron (ID: rnd-2_family-1) is distributed on chromosomes. We found that, in all chromosomes except chromosome 15, the Helitron subfamily sequences are accumulated at both ends of the chromosomes (Fig. 2a–c, S2, S3). In the regions of the local accumulation, the subfamily of Helitron (ID: rnd-2_family-1) showed tandem array with variety of unit lengths (Fig. 2b,c). We generated a self-sequence alignment of the consensus sequence of the subfamily of Helitron (ID: rnd-2_family-1) and found the repetitive nature of the consensus sequence (Fig. S3b). Copy numbers and genome coverage of the Helitron subfamily vary between chromosome arms and median copy number was148 copies and median genome coverage was 1.46 Mb (Fig. S3c,d). We also observed a tandem array of a single subfamily of an unclassified element (ID: rnd-1_family-0), beginning from the terminal of the array of a subfamily of Helitron (ID: rnd-2_family-1) and extending continuously to the end of the chromosome (Fig. 2a–c, S3a). Self-sequence alignment of the consensus sequence of unclassified element (ID: rnd-1_family-0) also revealed its repetitive nature (Fig. S3e). The copy numbers and genome coverage of the unclassified element (ID: rnd-1_family-0) differed across chromosome arms, with a median of 12,278 copies and a median genome coverage of 2.46 Mb (Fig. S3f,g). There was a statistically significant correlation between the copy number of the subfamily of Helitron (ID: rnd-2_family-1) in each chromosome arm and the copy number of the unclassified element (ID: rnd-1_family-0) in each chromosome arm (Fig. S3h), as well as between the genome coverage of the subfamily of Helitron (ID: rnd-2_family-1) in each chromosome arm and the genome coverage of the unclassified element (ID: rnd-1_family-0) in each chromosome arm (Fig. S3i). No statistically significant Blast hits were obtained between the consensus sequence of the subfamily of Helitron (ID: rnd-2_family-1) and that of the unclassified element (ID: rnd-1_family-0). We also found that the IS3EU subfamily (rnd-3_family-1290) were highly accumulated at the center of each chromosome (Fig. 2a,d, S2).

**Fig. 2.**
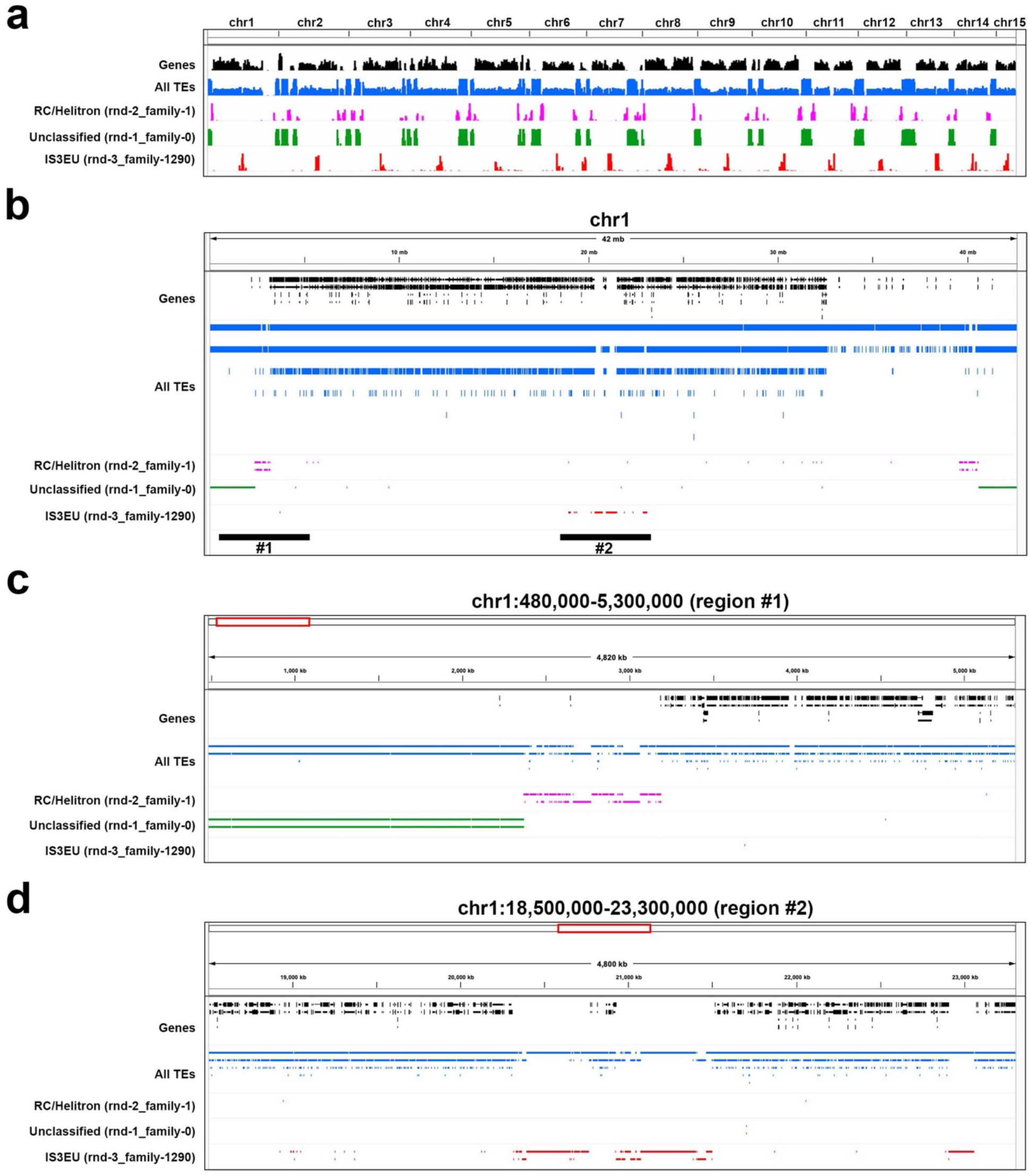
Genome localization of Helitron. (**a**) Whole genome landscape of Helitron localization. The first track indicates the positions of protein-coding genes, the second track represents all TEs, the third track represents the single subfamily of Helitrons labeled with rnd-2_family-1, and the fourth track depicts unclassified elements labeled with rnd-1_family-0. The fifth track shows the single IS3EU subfamily labeled with rnd-3_family-1290. (**b**) Helitron localization on the chromosome 1. The regions highlighted by black horizontal lines (#1 and #2) are further emphasized in panel (c) and (d). (**c**) The regional plot on the chromosome 1 (region #1). The first to fifth tracks are the same as in panels (a) and (b). (**d**) The regional plot on the chromosome 1 (region #2). The first to fifth tracks are the same as in panels (a) and (b).

For cross validation, we analyzed another genome assembly of *H. symbiolongicarpus* [35]. The contig-level genome assembly of *H. symbiolongicarpus* [35] was organized into 15 pseudochromosomal sequences using the publicly available female genetic linkage map [45]. Then, TE landscape was examined in these 15 pseudochromosomes. The subtelomeric accumulation of the subfamily of Helitron (ID: rnd-2_family-1) were also detected in the pseudochromosomes (Fig. S4a,b). The subtelomeric accumulation of the unclassified element (ID: rnd-1_family-0) and the centromeric accumulation of the IS3EU subfamily (rnd-3_family-1290) were not as distinct as in the chromosome-level genome assembly [37](Fig. S4a,b). This is most likely due to the nature of pseudochromosomes, where contigs were simply appended to female linkage groups. This could lead to the relatively low resolution in the subtelomeric and centromeric regions.

### Molecular phylogenetic analysis of Helitrons in the genome of *H. symbiolongicarpus*

Molecular phylogenetic analysis was conducted for copies of the Helitron subfamily (ID: rnd-2_family-1) and the unclassified element (ID: rnd-1_family-0) detected in the *H. symbiolongicarpus* genome in order to resolve their phylogenetic relationship. Examining the sequence length distribution of the 4,938 copies from the Helitron subfamily revealed a broad distribution, with a median length of 2,149 bp (Fig. 3a). In contrast, the 382,119 copies of the unclassified element (ID: rnd-1_family-0) exhibited a narrow length distribution, with a median length of 209 bp (Fig. 3b). From the Helitron subfamily, 177 sequences with lengths between 2.3 kb and 2.6 kb were extracted and aligned, resulting in a 3,419 bp alignment (Fig. 3c). Numerous insertions, deletions, and base substitutions were observed, suggesting that these sequences represent inactivated Helitrons (Fig. 3c). Similarly, 200 sequences of the unclassified element with lengths between 205 bp and 215 bp were randomly extracted and aligned, yielding comparable result (Fig. 3d). Furthermore, a molecular phylogenetic analysis using the neighbor-joining method was performed on the extracted sequences of the Helitron subfamily. This analysis identified monophyletic groups consisting of copies located on the same chromosome arm, suggesting their local propagation (Fig. 3e).

**Fig. 3.**
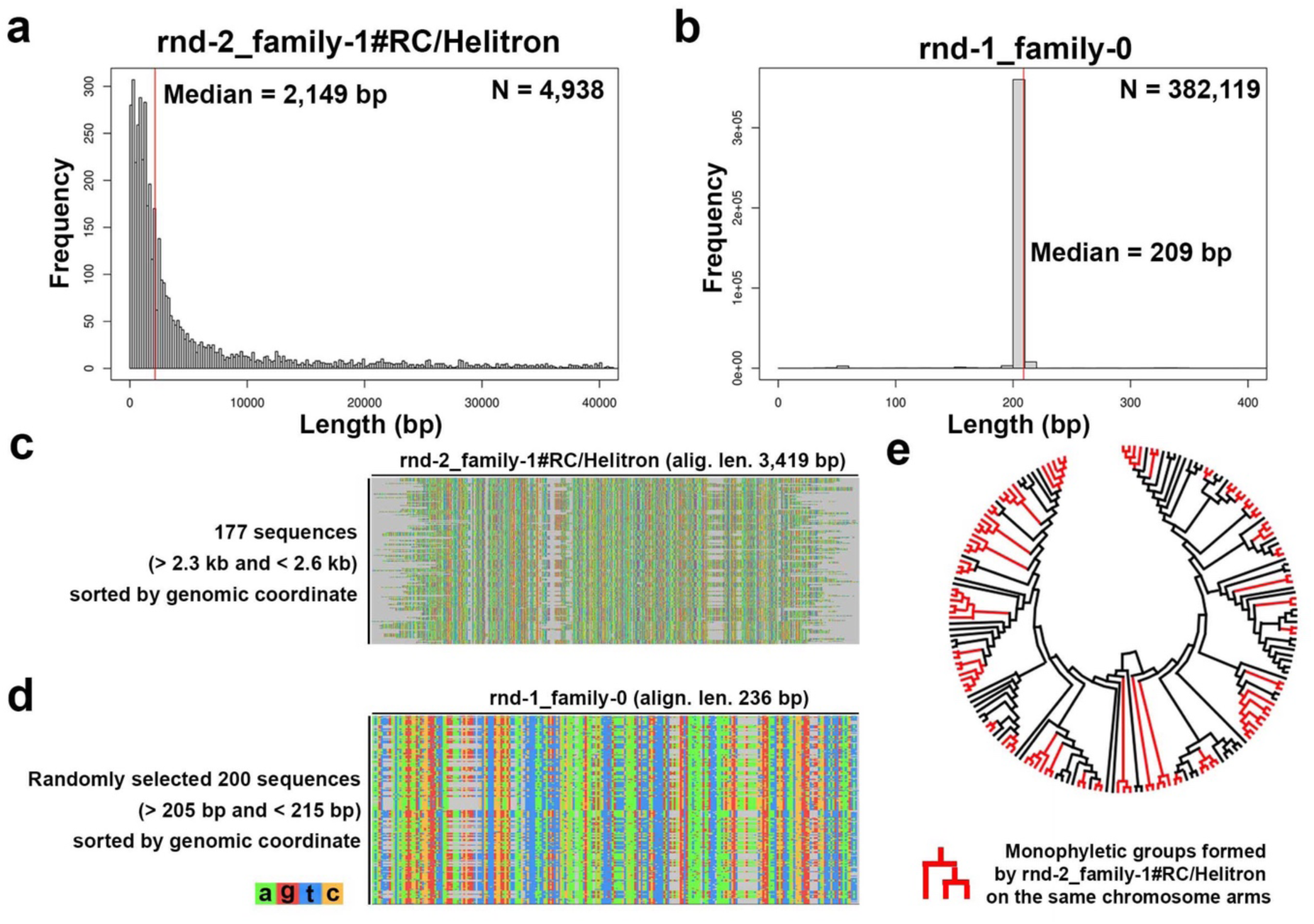
Molecular phylogenetic analysis of the Helitron subfamily in *H. symbiolongicarpus* genome. (**a**) Sequence length distribution of the Helitron subfamily labeled as rnd-2_family-1. (**b**) Sequence length distribution of the unclassified elements labeled as rnd-1_family-0. (**c**) Multiple sequence alignment of all 177 Helitron sequences labeled as rnd-2_family-1, with sequence lengths greater than 2.3 kb and less than 2.6 kb. (**d**) Multiple sequence alignment of 200 randomly selected unclassified sequences labeled as rnd-1_family-0, with sequence lengths greater than 205 bp and less than 215 bp. (**e**) Cladogram of a neighbor-joining tree of sequences from panel (c). Monophyletic groups formed by Helitron sequences from the same chromosome arms are highlighted in red.

## Genomic Distribution of Helitrons in Cnidaria

We investigated the presence of Helitron sequences in the genome assemblies of five additional cnidarian species (*Hydra vulgaris* [3], *Hydra viridissima* [38], *Rhopilema esculentum* [39], *Acropora millepora* [46], and *Nematostella vectensis* [40]). Helitrons were identified in all species analyzed (Fig. 4a). Notably, the genome coverage of Helitrons was higher in *N. vectensis* and *H. symbiolongicarpus* compared to the other species (Fig. 4a,b). To further explore their relationships, we extracted consensus sequences of Helitron subfamilies with lengths between 2 kb and 4 kb from each species and constructed a sequence alignment (Fig. 4c). The resulting alignment spanned 5,211 bp, suggesting that the Helitrons detected in the analyzed cnidarians are likely orthologous (Fig. 4c). Among the analyzed cnidarians, *N. vectensis*, similar to *H. symbiolongicarpus*, exhibited an accumulation of Helitrons near the chromosomal termini (Fig. 4d, upper panel). These regions were characterized by an extremely low gene density compared to other genomic regions (Fig. 4d, middle and bottom panels). In contrast, IS3EU elements, which were found to accumulate in the putative centromeric regions of *H. symbiolongicarpus* (Fig. 2, S2), displayed a diffuse distribution across the chromosomes in *N. vectensis* (Fig. 4d).

**Fig. 4.**
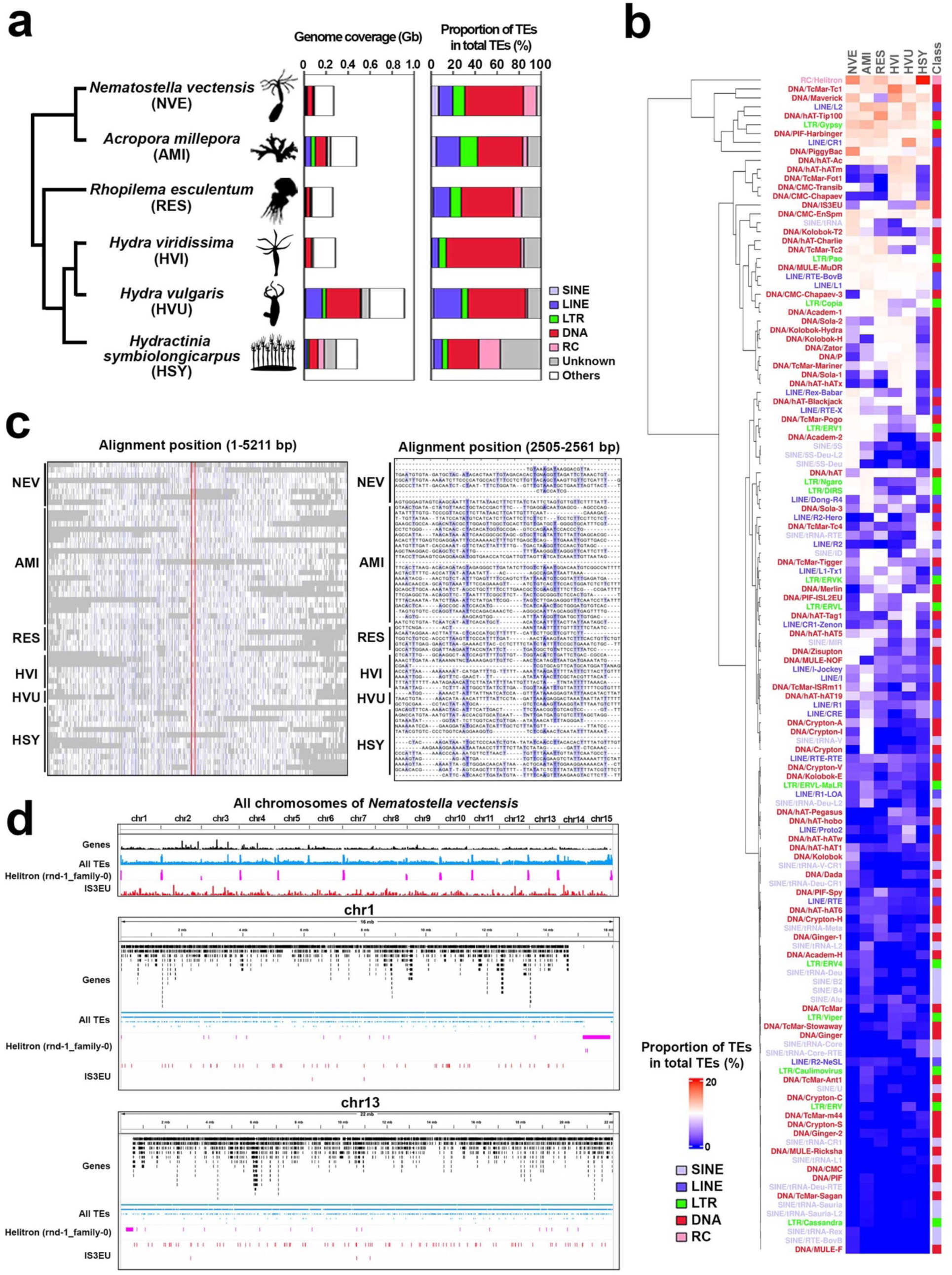
Helitrons in cnidarian genomes. (**a**) Cladogram of the six cnidarian species with their TE genome coverage and proportion of TEs in total TEs. (**b**) Proportion of TEs in total TEs across the six cnidarian species, with rows representing TE families, sorted by hierarchical clustering using the Euclidean distance and the Ward D2 methods. Columns represent species, all of which show varying amounts of Helitrons. (c) Sequence alignment of cnidarian Helitrons. The left panel shows a multiple sequence alignment of 58 Helitron consensus sequences with sequence lengths greater than 2 kb and less than 4 kb from six cnidarians. The right panel highlights the alignment between 2505 and 2561 bp. (d) Genomic localization of Helitrons in *Nematostella vectensis*, highlighting a subfamily (rnd-1_family-0) that is abundant in subtelomeric regions. Abbreviations of species names are as follows: NVE, *Nematostella vectensis*; AMI, *Acropora millepora*; RES, *Rhopilema esculentum*; HVI, *Hydra viridissima*; HVU, *Hydra vulgaris*; HSY, *Hydractinia symbiolongicarpus*.

## Discussion

Helitrons, or rolling-circle transposons, are a type of DNA transposon proposed to amplify through the rolling-circle replication (RCR) mechanism [11,12]. In the genomes of several plants, such as *Arabidopsis thaliana*, it has been reported that Helitrons form tandem arrays within the genome, which is probably a result of RCR generating tandem Helitron concatemers [47]. Additionally, in *Brassicaceae* species, Helitrons are found in lower densities in areas of the genome with high gene density, and they tend to preferentially insert into regions near centromeres and intergenic regions [48]. These findings have primarily been derived from analyses of plant genomes, leaving much to be explored regarding the role of RC transposons in metazoan genomes. In animal genomes, particularly in mammals, Helitrons seem to be less common and less active in comparison to plant genomes [12]. As an example of Helitron researches in mammals, in the bat *Myotis lucifugus*, Helitrons make up approximately 6% of the genome, and in several cases, they include one or two gene fragments [17]. Due to the sequence divergence of Helitrons, identifying them within repetitive element sequences requires highly sensitive detection methods[3]. Indeed, in our current study, running RepeatModeler with default parameters proved insufficient for detecting Helitrons in the *H. symbiolongicarpus* genome. With the modified RepeatModeler pipeline, we found the subtelomeric accumulations of a single Helitron subfamily in *H. symbiolongicarpus* and *N. vectensis*. By applying this approach to other animal genomes, we may be able to shed light on the potentially underestimated role that Helitrons could play in metazoan genomes, offering new insights into their evolutionary significance.

In this study, among the six analyzed cnidarian species (*H. symbiolongicarpus*, *Hydra vulgaris*, *Hydra viridissima*, *Rhopilema esculentum*, *Acropora millepora*, and *N. vectensis*), accumulations of IS3EU DNA elements were found in the putative centromeric regions of the *H. symbiolongicarpus* genome. DNA sequences of IS3EU DNA elements have been identified in the genomes of *Danio rerio* (zebrafish) and *Corbicula fluminea* (Asian clam) [49,50], and deposited in the RepBase database [51]. IS3EU is a DNA transposon belonging to the IS3 family [52]. Although multiple sequences of IS3EU are registered in Repbase Reports [49,50], detailed analyses focusing on their structure and function remain largely unexplored, leaving room for future investigation. Unlike Helitrons, no significant expansion of IS3EU was detected in *N. vectensis*, suggesting that the proliferation of IS3EU is specific to the *H. symbiolongicarpus* lineage.

The availability of a chromosome-scale genome assembly for *H. symbiolongicarpus* played a pivotal role in enabling a comprehensive characterization of the distribution of Helitrons within the genome. While it is feasible to analyze the average proportion of TEs across the genome using scaffold-or contig-level assemblies, this study revealed that Helitrons significantly enrich at the ends of chromosomes, contributing to an increase in genome size. Additionally, recent research on other TEs has shown that Ty3/Gypsy elements are highly overrepresented in pericentromeric and subtelomeric regions in frogs using a high-quality chromosome-scale genome assembly [53]. Such phenomena provide crucial insights into how TEs drive genome evolution, and detecting these patterns is greatly facilitated by high-quality chromosome-scale genome assemblies. Thorough investigations of chromosome-scale genome assemblies across various species could uncover novel TE distribution patterns that are dependent on specific genomic regions. This, in turn, may illuminate the role of TEs in maintaining chromosomal integrity, dynamics and evolution.

In this study, the accumulation of Helitrons was observed not only in the *H. symbiolongicarpus* genome but also in other cnidarian genomes such as the *N. vectensis* genome. *H. symbiolongicarpus* and *N. vectensis* belong to Medusozoa and Anthozoa, respectively, both of which are two major clades of Cnidaria that diverged over 500 million years ago [54,55]. Building on these findings, our study suggests the significant evolutionary role of Helitrons in cnidarian genomes, particularly in shaping the telomeric structure. The conserved chromosomal-level homology observed across cnidarian species, coupled with the accumulation of Helitrons in distinct genomic regions of both medusozoa and anthozoa lineages, suggests the potential influence of Helitrons in maintaining genomic stability over vast evolutionary timescales. As high-quality chromosomal assemblies continue to become available for cnidarian species [3,37,39,56], further exploration of TE dynamics across diverse lineages will help identify the ancestral chromosomal characteristics. Together with other eukaryotic studies on Helitron distribution [20] this will shed light on broader patterns of Helitron-driven genomic innovation in metazoans.

## Methods

### Generation of custom repetitive element libraries and TE detection in genomes

Annotation of TEs in genomes can sometimes be challenging, largely depending on the extent to which the transposons of the target species or its close relatives are represented in known databases such as Dfam [59]. This difficulty arises because most TEs become inactivated and degraded over time due to the accumulation of mutations, including base substitutions, insertions, and deletions. [41,42]. To overcome this challenge, we adopted an approach similar to that used in a previous study[3]. First, we obtained the RefSeq hydractinia genome sequence from the NCBI Genome database (NCBI RefSeq accession GCF_029227915.1) [37] and generated a custom repeatitive element library using RepeatModeler v 2.0.5 [43] with the sequence as the query. Next, we downloaded all Dfam HMM profiles of TEs from the Dfam database (https://dfam.org/home). Due to the large size of the Dfam HMM profile dataset for TEs, we divided it into smaller chunks, each containing 25,000 HMM profiles. Each chunk file was indexed using hmmpress [44]. We then used the sequences of the custom repeat library as queries to perform a HMM-based search against the Dfam HMM profiles using nhmmscan with default parameters [44]. For each sequence in the custom repeatitive element library, we identified the hit sequences in the Dfam database with the lowest E-value, as long as it was less than 0.05. Thereby we refined the annotations in the custom repeat library based on the search results. Using the refined custom repetitive element library, we ran RepeatMasker [60] on the genome sequences with the options of -parallel 70 -gff -a -dir - xsmall. For *Hydra vulgaris* (GCA_029227915.2), *Hydra viridissima* (GCA_014706445.1), *Rhopilema esculentum* [39], *Acropora millepora (GCF_013753865.1)*, and *N. vectensis (GCF_932526225.1),* we retrieved genome sequences from the NCBI Genome database and performed repetitive element analyses in the same manner as for *H. symbiolongicarpus*. We extracted TE regions from the RepeatMasker output files and analyzed TE distributions. Gene annotation of each species was retrieved from the NCBI Genome database. The positions of genes and TEs on the genomes were visualized using Integrative Genomics Viewer (IGV) [61].

### Analysis using an alternative genome assembly of *H. symbiolongicarpus*

An alternative genome assembly of *H. symbiolongicarpus* (Hsym_primary_v1.0 [35]) was obtained from the Hydractinia Genome Project Portal (https://research.nhgri.nih.gov/hydractinia/). Because the Hsym_primary_v1.0 is a conti-level genome assembly, we generated a pseudo-chromosomal sequences by anchoring the contig sequences of the Hsym_primary_v1.0 assembly to the previously reported female genetic linkage maps of *H. symbiolongicarpus* [45]. In this process, we aligned the contig sequences based on the SNP marker positions [45] and constructed 15 pseudochromosome sequences. The TE landscape on the 15 pseudochromosome sequences was analyzed using RepeatMasker with a custom repetitive element library generated from the RefSeq Hydractinia genome assembly (NCBI RefSeq accession GCF_029227915.1) following the method described above.

### TE Distribution Analyses

For the analysis of Kimura substitution levels, we extracted the Kimura substitution values for each TE from the RepeatMasker output file and visualized them using the barplot function in R (https://www.R-project.org/.). Similarly, the genome-wide coverage of TEs was calculated based on data extracted from the RepeatMasker result. The proportions of various TE contents were visualized as a heatmap using the ComplexHeatmap R library [62]. For generating the self-alignments of consensus sequences of TE families, alignments were generated using blastn [63], followed by visualization with the blast2dotplot.py script from the bio_small_scripts repository (https://github.com/satoshikawato/bio_small_scripts). DNA sequence multiple alignments were generated using Clustal Omega [64] with its default parameters. The resulting multiple sequence alignments were visualized using Jalview [65]. Molecular phylogenetic analyses were performed using MEGA software [66].

## Data availability

This study did not involve the generation of new sequencing data. All analyses were conducted using publicly available datasets, as detailed in the Methods section, which served as the primary data sources for this research.

## Acknowledgments

The authors thank Uri Frank at the University of Galway for comments on the manuscript, as well as Nina Znidaric, Angela Caballero Alfonso, and Eduard Renfer at the University of Vienna for their technical assistance. The computation for this study was performed at the Life Science Compute Cluster (LiSC) of the University of Vienna.

## Author contributions

TK, KK, and OS designed the study. TK, KK, and OS analyzed the data. TK, KK, and OS wrote the manuscript. All authors reviewed the manuscript and agreed to the content.

## Supplementary figures

**Fig. S1.**
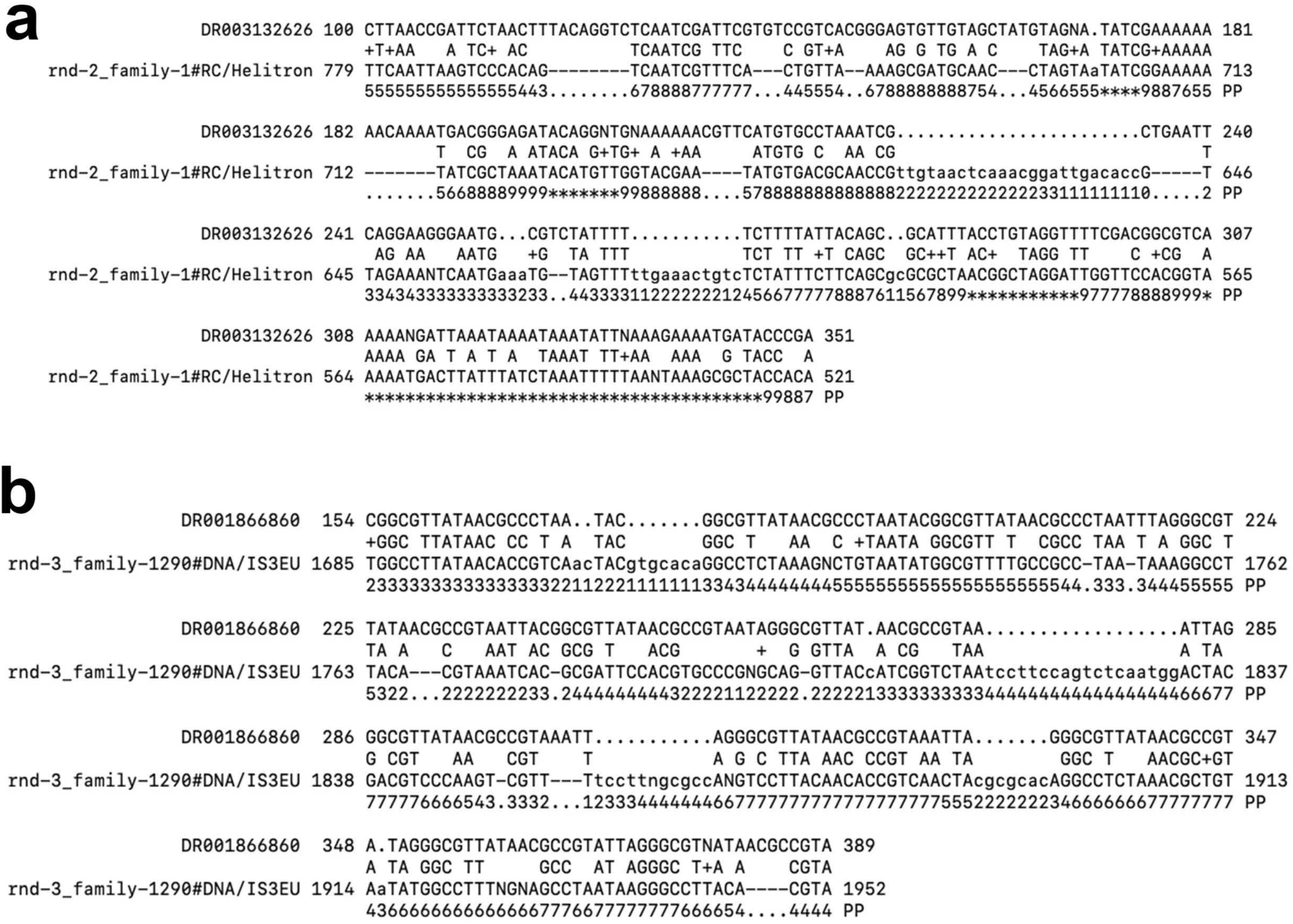
TE subfamily consensus sequences of *H. symbiolongicarpus* from the custom repetitive element library and their homologous sequences in the Dfam database. **(a)** Detected homologous sites between the consensus of the Helitron subfamily (rnd-2_family-1) and the Helitron sequence (Dfam accession number DR003132626). **(b)** Detected homologous sites between the consensus sequence of the IS3EU subfamily (rnd-3_family-1290) and the IS3EU sequence (Dfam accession number DR001866860).

**Fig. S2.**
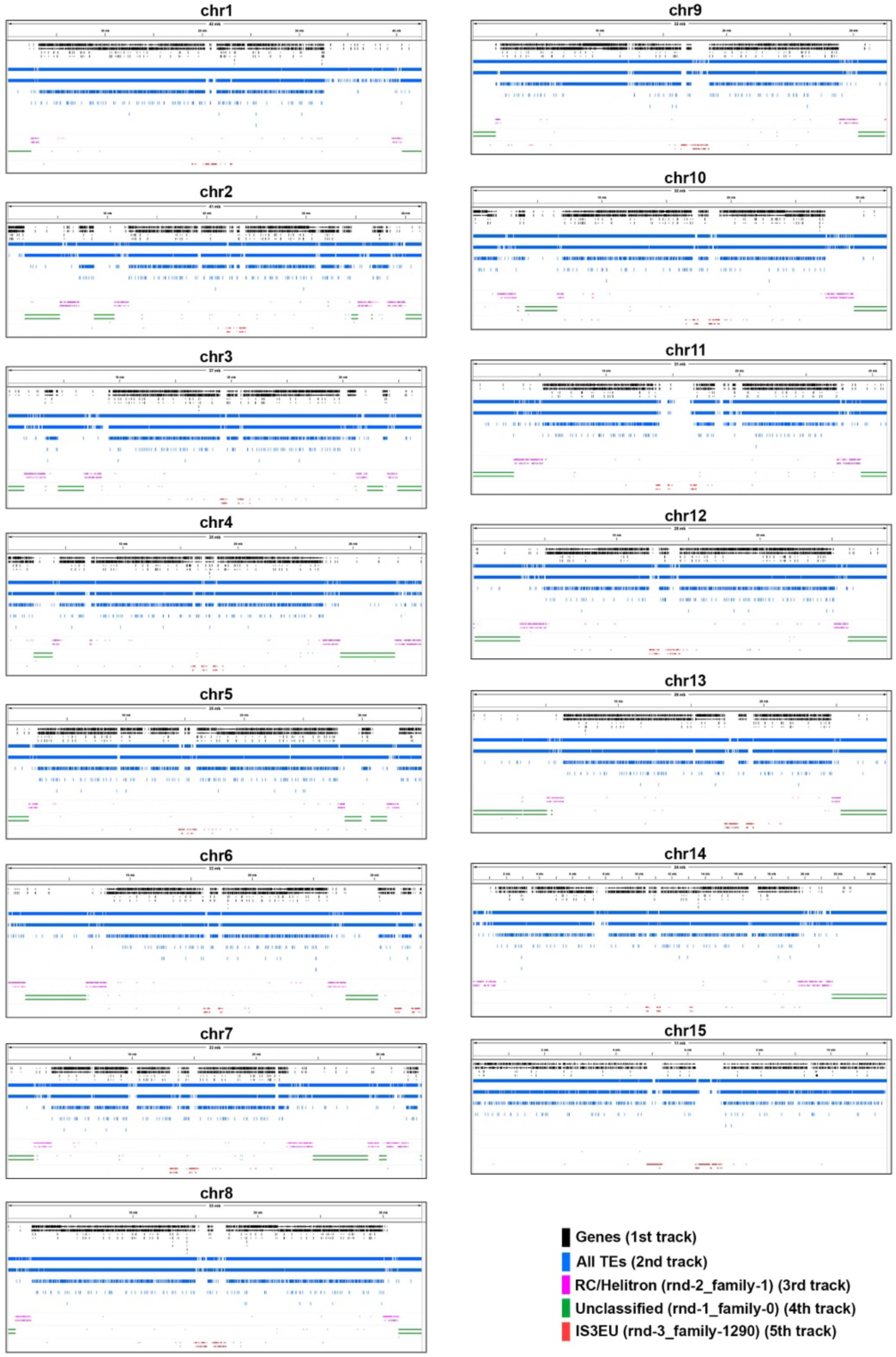
Helitron localization in each chromosome. The first track shows the positions of protein-coding genes, the second track represents all TEs, the third track represents the single subfamily of Helitrons labeled with rnd-2_family-1, and the fourth track depicts unclassified elements labeled with rnd-1_family-0. The fifth track shows the single IS3EU subfamily labeled with rnd-3_family-1290.

**Fig. S3.**
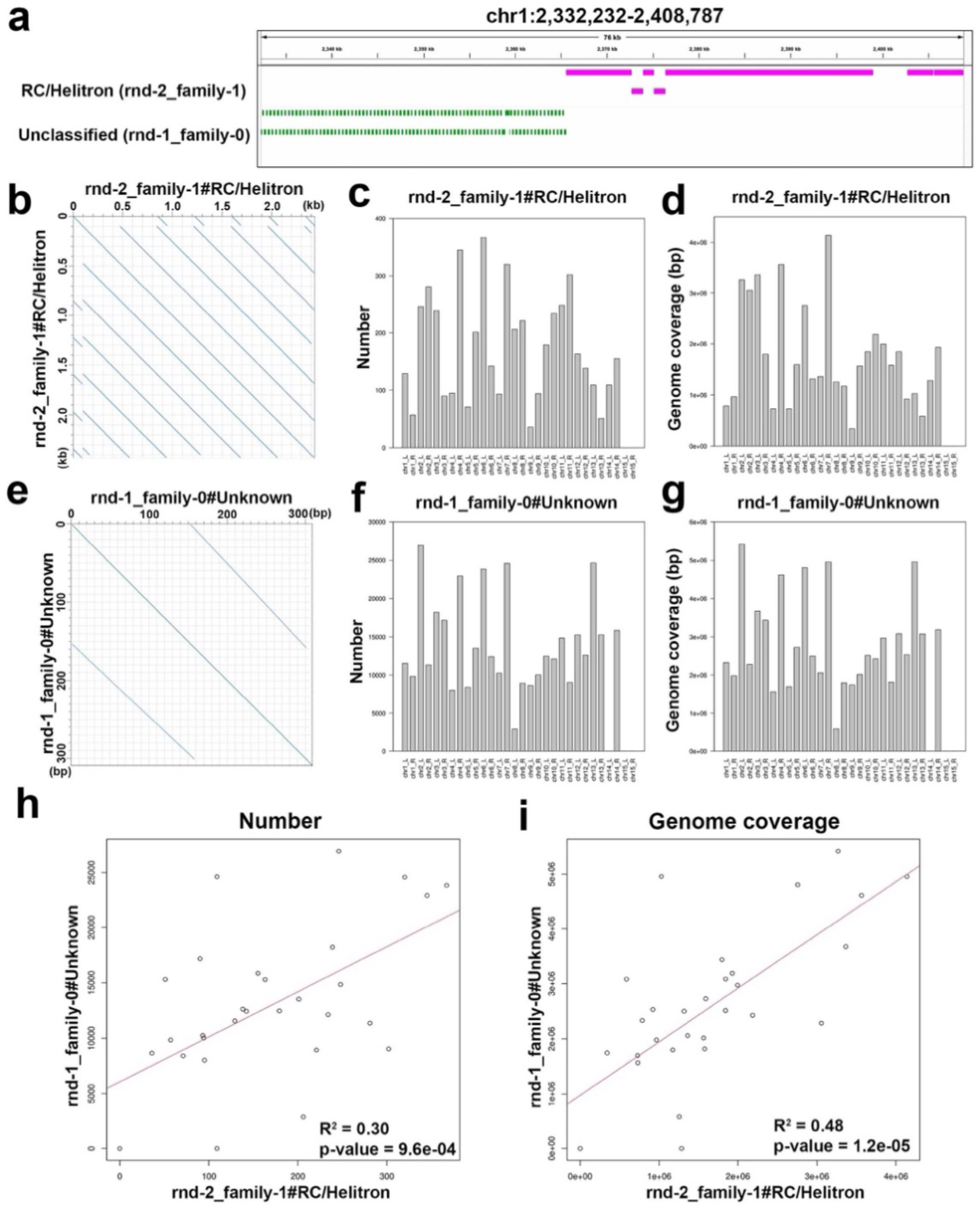
Copy numbers and genome coverage of the single Helitron family. **(a)** Regional distribution of the subfamily of Helitron (rnd-2_family-1) and the unclassified subfamilies (rnd-1_family-0) on chromosome 1 base from 2,326,326 bp to 2,402,881 bp. **(b)** Self-sequence alignment of the consensus sequence of the subfamily of Helitron (rnd-2_family-1). **(c)** Number of the subfamily of Helitron (rnd-2_family-1) in each chromosome arm. **(d)** Genome coverage of the unclassified subfamily of Helitron (rnd-2_family-1) in each chromosome arm. **(e)** Self-sequence alignment of the consensus sequence of the unclassified subfamily (rnd-1_family-0). **(f)** Number of the unclassified subfamily (rnd-1_family-0) in each chromosome arm. **(g)** Genome coverage of the unclassified subfamily (rnd-1_family-0) in each chromosome arm. **(h)** Correlation analysis between number of the subfamily of Helitron (rnd-2_family-1) and that of the unclassified subfamilies (rnd-1_family-0). **(i)** Correlation analysis between genome coverage of the subfamily of Helitron (rnd-2_family-1) and that of the unclassified subfamilies (rnd-1_family-0).

**Fig. S4.**
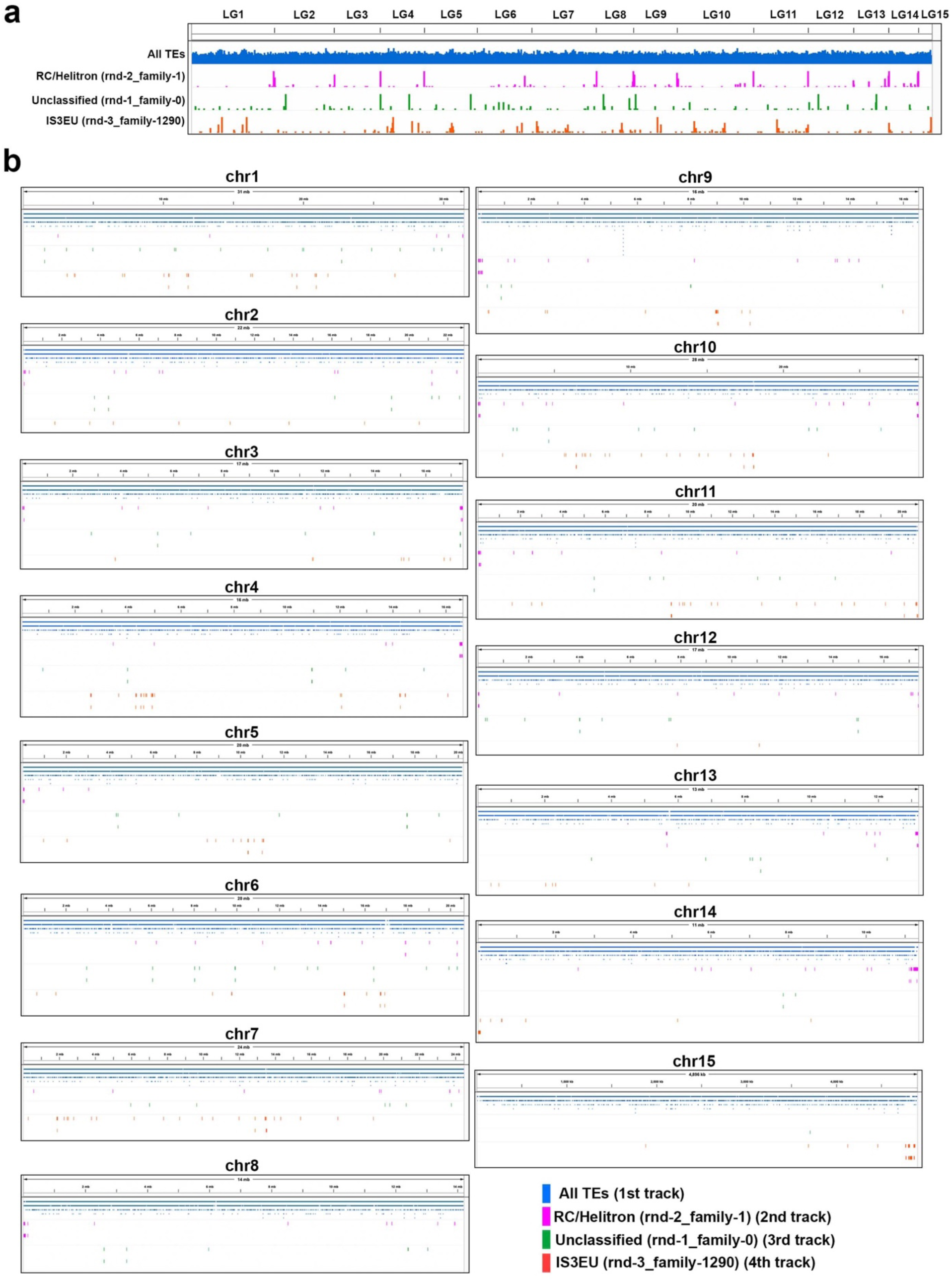
Helitron localization on the pseudochromosomes generated by the Hsym_primary_v1.0 assembly and the maternal genetic linkage map. (a) Helitron distribution across all pseudochromosomes. Pseudochromosomes were generated by the Hsym_primary_v1.0 assembly [35] and the maternal genetic linkage map [45]. Note the accumulation of the Helitron subfamily (rnd-2_family-1) at the ends of chromosomes. (b) Helitron distribution in each pseudochromosomes. The track arrangement is the same as in panel (a).

## Supplementary Tables

**Table S1. Genome coverage of major TE groups**

**Table S2. Genome coverage of TE families**

**Table S3. Genome coverage of TE subfamilies**

**Table S4. List of cnidarian species analyzed in this study**

## Abbreviations

TEs: Transposable elements
RC: rolling-circle
RCR: rolling-circle replication
HMM: Hidden Markov model
HSY: *Hydractinia symbiolongicarpus*.
HVU: *Hydra vulgaris*
HVI: *Hydra viridissima*
RES: *Rhopilema esculentum*
AMI: *Acropora millepora*
NVE: *Nematostella vectensis*

